# miRNA-encoded regulatory peptides modulate cadmium tolerance and accumulation in rice

**DOI:** 10.1101/2023.07.22.550150

**Authors:** Long Lu, Xinyu Chen, Jiaming Chen, Zaoli Zhang, Zhen Zhang, Yanyan Sun, Yuan Wang, Siwen Xie, Yinuo Ma, Yuanyuan Song, Rensen Zeng

**Author notes:** ^1^ These authors contributed equally to this work (Long Lu, Xinyu Chen). Corresponding Authors Rensen Zeng Yuanyuan Song.

## Abstract

MicroRNAs (miRNAs) are small non-coding RNAs that play a vital role in plant responses to abiotic and biotic stress. Recently, it has been discovered that some primary miRNAs (pri-miRNAs) encode regulatory short peptides called miPEPs. However, the presence of miPEPs in rice, and their functions in response to abiotic stress, particularly stress induced by heavy metals, remain poorly understood. Here, we identified a functional small peptide (miPEP156e) encoded by pri-miR156e that regulates the expression of miR156 and its target *SPL* genes, thereby affecting miR156-mediated cadmium (Cd) tolerance in rice. Overexpression of *miPEP156e* led to a reduction in the accumulation of reactive oxygen species (ROS) and Cd in plants under Cd stress, resulting in improved rice Cd tolerance, as observed in miR156-overexpressing lines and seedlings treated with exogenous miPEP156e. In contrast, *miPEP156e* mutants displayed sensitivity to Cd stress due to the elevated accumulation of ROS and Cd. Transcriptome analysis further revealed that miPEP156e improved rice Cd tolerance by modulating Cd transporter and ROS scavenging genes. Moreover, we identified five novel miPEPs involved in regulating Cd resistance through exogenous treatment of seedlings with synthetic corresponding miPEPs. Our study provides insights into the regulatory mechanism of miPEP156e in rice response to Cd stress and demonstrates the potential of miPEPs as an effective tool for improving crop abiotic stress tolerance.

## Introduction

Toxic mineral elements such as cadmium (Cd) and lead (Pb) are widespread in the Earth’s crust. Among the toxic minerals, heavy metal Cd is highly toxic to both humans and plants and is non-essential for plant growth and development. Human activities are strongly associated with excessive levels of Cd in water and soil (Song et al., 2017). Over-accumulated Cd in plants can cause heavy metal toxicity, disrupt nutrient and reactive oxygen species (ROS) homeostasis, interfere with photosynthesis, and threaten crop production and food safety (Zhao and Wang, 2020). Cd is easily absorbed by plants and can be translocated and accumulated in edible parts, such as grains and leaves (Chang et al., 2020). As the main staple food worldwide and a major source of human exposure to Cd (Shimbo et al., 2001), rice’s uptake and translocation of Cd need to be reduced to prevent Cd from entering the food chain.

To alleviate the damage caused by Cd, plants have evolved a variety of strategies to cope with Cd toxicity, including microRNA (miRNA)-directed gene expression regulation at the post-transcriptional level (Carrington and Ambros, 2003; Zhou et al., 2012; Tang et al., 2014). In the past decade, numerous Cd-responsive miRNAs and their target genes have been identified through transcriptome and small RNA sequencing technology in *Arabidopsis*, rice, and other species (Fang et al., 2013; Fu et al., 2019; Gao et al., 2019; Zhou et al., 2019). Cd uptake and transport in plants are mediated by the transporters responsible for Fe, Zn, or Mn, such as ATPase binding cassette transporter (ABC) and natural resistance-associated macrophage protein (NRAMP) (Park et al., 2012; Wu et al., 2016; Chang et al., 2020). Among the Cd-responsive miRNA in rice, miR192 and miR268 target genes encoding ABC and NRAMP transporter proteins, respectively, affecting seedling growth (Ding et al., 2013; Ding et al., 2017). Approximately 66% of verified miRNAs targets are transcription factors (TFs), implying these TFs may be critical regulators of plant responses to heavy metal stress (Tang and Chu, 2017). For instance, the *HOMEODOMAIN CONTAINING PROTEIN 4* (*OsHB4*) gene, a target of miR166, plays a negative role in Cd tolerance and accumulation in rice (Ding et al., 2018).

miR156, a highly conserved miRNA in land plants, was first discovered to be a master regulator of vegetative phase transition in *Arabidopsis*, its regulatory role has subsequently been widely observed in other species (Wang et al., 2009; Wu et al., 2009; Xie et al., 2012; Liu et al., 2017). Several studies have shown that miR156 and its targets SQUAMOSA-promoter binding protein-like (SPL) TFs function in the regulation of crop productivity and stress tolerance (Chen et al., 2015; Ge et al., 2018; Miao et al., 2019). In rice, microarray analysis and high-throughput sequencing have revealed that miR156 is responsive to Cd stress (Ding et al., 2011; Zhong et al., 2019). However, the role of miR156 in Cd accumulation and tolerance in rice remains unknown.

Although it is generally assumed that non-coding RNAs (nc-RNAs) do not encode peptides, recent studies have shown that short open reading frames (ORFs) in some pri-miRNAs can code for short peptides called miPEPs, which have been found in both animals and plants (Lauressergues et al., 2015; Kang et al., 2020; Lauressergues et al., 2022). These miPEPs influence the accumulation of the associated miRNAs and hence regulate the expression of target genes. Several miPEPs have been characterized and found to regulate plant growth and development, some of which are considered novel and effective tools for crop improvement. For example, the application of synthetic miPEP172c increased nodulation in soybean by enhancing the activity of miR172c (Couzigou et al., 2016). This was consistent with a previous study, which demonstrated that overexpression of miR172c in soybean increased nodule number (Wang et al., 2014). Another recent study revealed that the primary transcript of miR171d encodes a regulatory peptide, vvi-miPEP171d1, which controls adventitious root formation in grapevine (Chen et al., 2020). To date, studies on miPEPs have mainly been conducted in dicotyledons, and it is still unknown whether the function of miPEPs is conserved in monocots, especially in important crops, such as rice, maize, and wheat.

In this study, we discovered a functional regulatory short peptide (miPEP156e) present in the primary transcript of miR156e, which had the highest induction by Cd treatment among miR156 family members. Application of synthetic miPEP156e resulted in the increased miR156 expression and downregulation of its target *SPL* genes. More interestingly, plants treated with miPEP156e and miR156-OE lines showed increased tolerance against Cd stress. To better understand the role of miPEP156e, we developed *miPEP156e*-OE lines and *miPEP156e*-Cr mutants. Similar to the miR156-OE lines, *miPEP156e*-OE plants accumulated less Cd and ROS by modulating the expression of genes encoding key Cd transporters and ROS scavenging genes, thus exhibiting enhanced Cd tolerance, while a contrasting phenotype was observed in *miPEP156e*-Cr mutants. Our results demonstrated the significance of miPEP156e in regulating miR156 activity and its associated Cd tolerance in rice.

## Results

### Expression profiles of *MIR156* family members in response to Cd stress

In rice, several Cd-responsive miRNAs, including miR528, miR171, miR166, miR156, and miR396, have previously been identified using microarray analysis and high-throughput sequencing (Ding et al., 2011; Tang et al., 2014; Zhong et al., 2019). However, only a few of these miRNAs have been characterized (Ding et al., 2013; Ding et al., 2016; Ding et al., 2017; Ding et al., 2018). To investigate the role of miR156 in Cd response, we analyzed the expression patterns of *MIR156* genes in rice roots under Cd stress by detecting the abundance of corresponding precursors. The results showed that the expression patterns varied among different members in response to Cd stress. The transcript levels of pre-miR156b, pre-miR156c, and pre-miR156l dynamically decreased upon Cd stress (Figure 1). Most of the other members were induced by Cd at 3 or 6 h (Figure 1). Among them, pre-miR156e exhibited the highest induction level (Figure 1), suggesting that *MIR156e* might play a crucial role in rice Cd tolerance.

**Figure 1.**
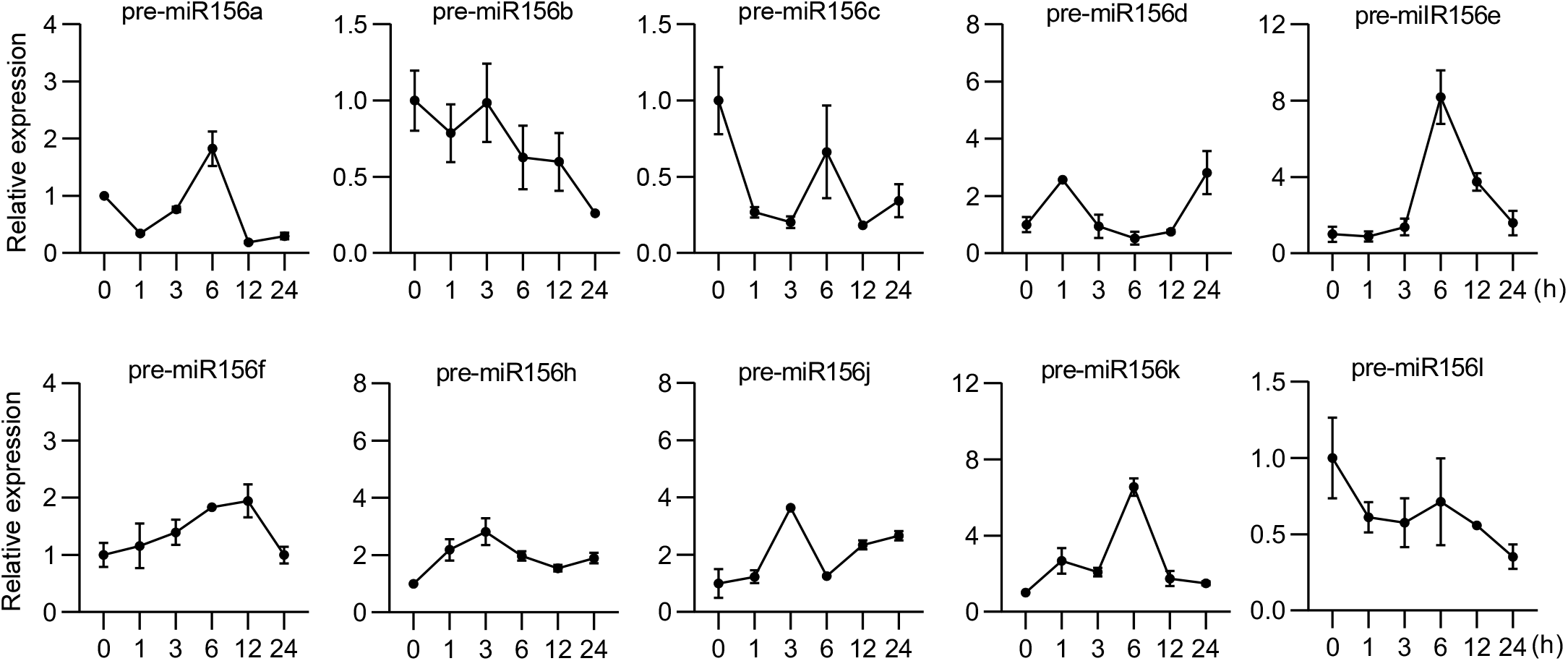
Expression of miR156 family members in response to Cd stress in rice. The results are presented as means ± SD of three biological replicates.

### Identification of miPEP156e in the primary transcript of miR156e

Since miPEPs modulate the accumulation of associated mature miRNAs by promoting pri-miRNAs expression, we speculated that there might be a miPEP in rice that specifically regulates the abundance of miR156e like other species, such as *Arabidopsis* and soybean. We designed different forward primers in the upstream regions of pre- miR156e and a reverse primer that targeted pre-miR156e for PCR amplification. The result showed that the 520 base pair (bp) sequence upstream of pre-miR156e was part of the primary transcript of miR156e (Supplemental Figure 1). Further analysis of the sequence led to the identification of four putative ORFs, encoding 36 (ORF1), 37 (ORF2), 27 (ORF3), and 11 (ORF4) amino acids, respectively. The translation initiation sites of these ORFs were named ATG1, ATG2, ATG3, and ATG4, respectively (Figure 2A). To determine the activity of these ORFs in planta, the translation initiation sites (ATG1, ATG3, and ATG4), along with the upstream promoter sequences, were fused to the *GUS* reporter gene, and the CaMV 35S promoter was used to drive the *GUS* gene as the positive control. The GUS activity was analyzed in *Nicotiana benthamiana* leaves. Although all three translation initiation sites (ATG1, ATG3, and ATG4) could initiate the transcription of the *GUS* gene, GUS staining analysis showed that the activity of the GUS protein was only detected when the *GUS* gene was fused to ATG1 (Figure 2B and 2C), indicating that the ATG1 of ORF1 was capable to initiate translation.

**Figure 2.**
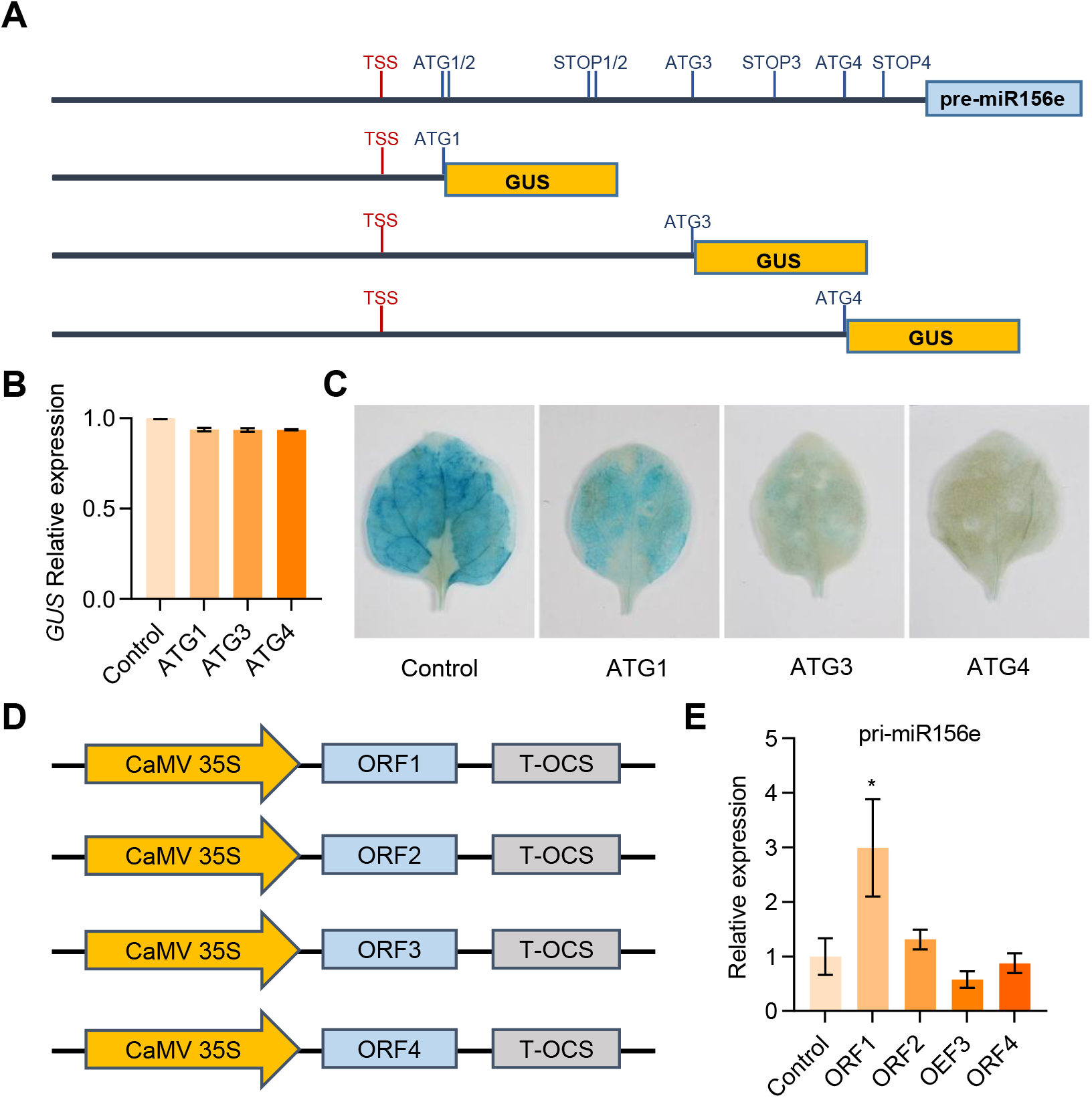
Identification of miPEP156e in the primary transcript of miR156e. A, Schematic diagrams of different constructs used for GUS assay in *Nicotiana benthamiana* leaves. TSS, transcription start site. ATG, translation initiation site. STOP, translation termination site. B, Expression of the *GUS* gene in *N*. *benthamiana* leaves. Bars indicate SD (n = 3). C, Histochemical staining showing GUS activity in *N*. *benthamiana* leaves transformed with different constructs. D, Schematic diagrams of the *35S_pro_*:*ORF1/2/3/4* constructs used for transient expression assay in rice protoplasts. E, Expression of pri-miR156e in rice protoplasts after transient expression of ORF1, ORF2, ORF3, or ORF4. Bars indicate SD (n = 3). Asterisks indicate significant differences compared with the corresponding controls (*, *P* < 0,05; **, *P* < 0.01).

We then used rice protoplast transient expression system to study the effect of four ORFs on the transcription of pri-miR156e, using the *GFP* gene under the control of CaMV 35S promoter as a control (Figure 2D). Real-time quantitative PCR (RT-qPCR) analysis showed that ORF1 promoted the transcript level of pri-miR156e, while the other three ORFs did not have this effect (Figure 2E). These results suggested that ORF1 was capable of regulating miR156e expression. Therefore, we named the short peptide encoded by ORF1 as miPEP156e.

### Effect of miPEP156 on the phenotype of rice seedlings under Cd stress

To explore whether synthetic miPEP156e can be absorbed by rice roots after the external application, we used the Fitc (fluorescein isothiocyanate)-labeled miPEP156e. Our results showed that 24 hours after applying Fitc-miPEP156e, the labeled peptide penetrated the root cap, and the fluorescence signal was observed throughout the root cells 48 hours after treatment (Figure 3A). We then used synthetic miPEP156e to examine its effect on Cd tolerance in rice. Under normal conditions, exogenous miPEP156e did not affect the growth of 19-day-old hydroponic seedlings. However, when exposed to Cd stress, the application of miPEP156e reduced seedling sensitivity to Cd stress (Figure 3B), resulting in a dose-response increase in root length and biomass after Cd treatment (Figure 3C and 3D). Similar results were obtained from an assay on 8-day-old seedlings cultured in paper pouches (Supplemental Figure 2). A concentration of 1 μM miPEP156e was selected as the optimal for further studies.

**Figure 3.**
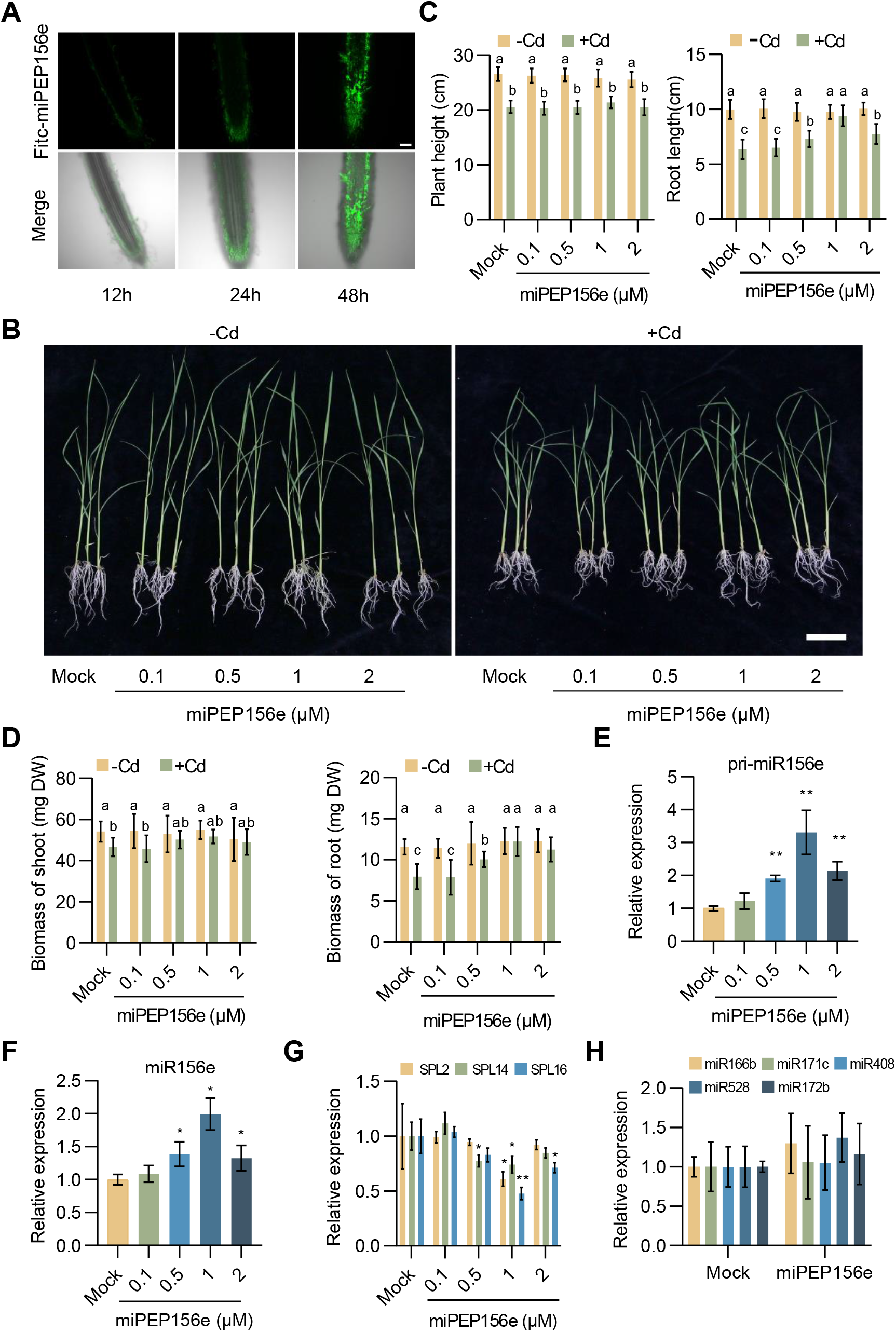
Influence of miPEP156e on miR156e expression and phenotypic changes of rice seedlings under Cd toxicity. A, Confocol images showing the uptake of 5 μM Fitc-miPEP156e in Nip roots after incubation for 12, 24, and 48 hours. Scale bar = 100 μm. B, Performance of 19-day-old hydroponic rice seedlings treated with water or different concentrations of miPEP156e in the presence of Cd stress. Scale bar = 5 cm. C, Plant height and root length of rice seedlings treated with water or different concentrations of miPEP156e under Cd stress conditions. Error bars indicate SD (n = 8). D, Biomass of shoots and roots of rice seedlings treated with water or different concentrations of miPEP156e under Cd stress conditions. Error bars indicate SD (n = 8). E, Quantification of pri-miR156e in the seedlings treated for 6h with water or synthetic miPEP156e. Error bars indicate SD (n = 3). F, G, Quantification of miR156e (F) and its target genes (*SPL2*, *SPL14*, and *SPL16*) (G) in the seedlings treated for 6h with water or synthetic miPEP156e. Error bars indicate SD (n = 3). H, Effect of exogenous treatment with miPEP156e on the expression of various miRNAs. Error bars indicate SD (n = 3). For C and D, the lowercase letters ‘a’ to ‘c’ indicate significant differences according to Tukey’s test (*P* < 0.05 or *P* < 0.01). Asterisks indicate significant differences compared with the corresponding controls (*, *P* < 0,05; **, *P* < 0.01).

Next, we investigated the effect of miPEP156e on the expression of miR156e and its target *SPL* genes. Exogenous miPEP156e significantly enhanced the transcript levels of pri-miR156e and miR156e, while repressing the expression of *SPL2*, *SPL14*, and *SP16* (Figure 3E, 3F, and 3G). The expression level increased or decreased with treatment duration. To study the specificity of miPEP156e, 19-day-old hydroponic seedlings were treated with ORF2-encoded short peptide (miPEP156e-ORF2). However, unlike miPEP156e, the application of miPEP156e-ORF2 did not affect the phenotype of rice seedlings in the presence of Cd stress and the expression of miR156e and its targets (Supplemental Figure 3).

In addition, to study whether miPEP156e specifically regulates miR156e, the response of several other miRNAs to miPEP156e was analyzed. Exogenous miPEP156e did not affect the expression of other miRNAs (Figure 3H), indicating that miPEP156e can improve rice tolerance to Cd stress and specifically modulate the abundance of miR156e and its targets.

### miR156 alleviates Cd toxicity-induced damage in rice

As miPEPs exert their function by affecting corresponding mature miRNA, we hypothesized that miR156 plays an essential role in rice Cd tolerance. Two individual miR156-overexpressing transgenic lines miR156-OE2 and miR156-OE5 (Liu et al., 2019), which showed reduced expression of the *SPL* genes (Supplemental Figure 4), were used to test Cd tolerance. Both wild-type ZH11 plants and transgenic plants grew well under normal conditions (Figure 4A). However, after Cd treatment, transgenic plants performed better than wild-type plants, as revealed by the longer roots and the higher plants (Figure 4A and 4B). Consistently, the biomass of roots and shoots was significantly higher in miR156-OE plants compared to wild-type ZH11 plants (Figure 4C). These results indicated that overexpression of miR156 confers Cd tolerance in rice.

**Figure 4.**
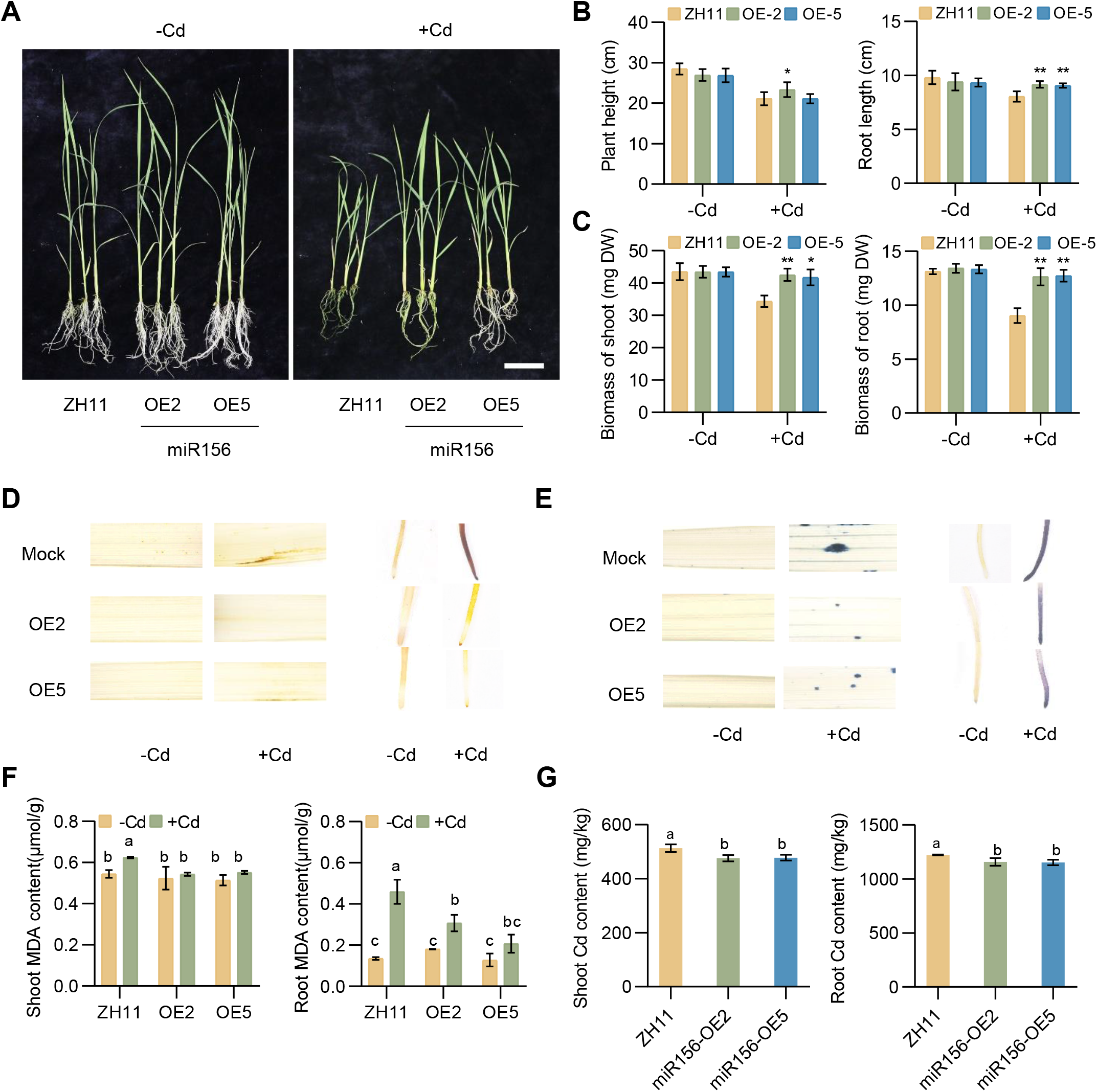
miR156 positively regulates Cd tolerance in rice. A, Overexpression of miR156 improved the performance of transgenic seedlings under Cd toxicity. B, Plant height and root length of wild-type and miR156-OE seedlings after Cd treatment. Error bars indicate SD (n = 8). C, Biomass of shoots and roots of wild-type and miR156-OE seedlings after Cd treatment. Error bars indicate SD (n = 8). D, DAB staining of leaves and roots of wild-type and miR156-OE seedlings after water or Cd treatment. The brown color indicates H_2O2 l_evel in each leaf and root. E, NBT staining of leaves and roots of wild-type and miR156-OE seedlings after water or Cd treatment. The blue color indicates O^2-^ level in each leaf and root. F, Measurement of MDA contents in shoots and roots of wild-type and miR156-OE seedlings. Error bars indicate SD (n = 3). G, Measurement of Cd contents in shoots and roots of wild-type and miR156-OE seedlings. Error bars indicate SD (n = 3). The lowercase letters ‘a’ to ‘c’ indicate significant differences according to Tukey’s test (*P* < 0.05 or *P* < 0.01).

Excessive accumulation of heavy metals can cause toxicity and induce high levels of reactive oxygen species (ROS) in plant cells, which can damage proteins, lipids, and other cellular components (Hu et al., 2020). Since miR156-OE plants displayed insensibility to Cd toxicity, we wondered whether the ROS accumulation might also be affected. The ROS levels in roots and leaves were analyzed through DAB and NBT staining to confirm this hypothesis. DAB staining revealed that miR156-OE plants accumulated less hydrogen peroxide (H_2_O_2)_ than wild-type plants (Figure 4D). Similarly, a high level of superoxide anion (O^2-^) was detected in wild-type plants upon Cd treatment by NBT staining, whereas a relatively lower accumulation of O^2-^ was observed in miR156-OE plants (Figure 4E). The malondialdehyde (MDA) and Cd content assay of wild-type plants and miR156-OE lines (Figure 4F and 4G) also showed similar results, indicating that miR156-OE plants have stronger antioxidant and detoxification abilities.

### Overexpression of *miPEP156e* regulates miR156e-associated phenotype

To determine the function of miPEP156e in regulating Cd resistance in rice, stable transgenic rice plants overexpressing *miPEP156e* driven by the CaMV 35S promoter were generated in the Nip background (Figure 5A). Three homozygous *miPEP156e*-overexpressing transgenic lines were obtained for further studies (Figure 5B). We first investigated whether *miPEP156e* overexpression could affect the expression of miR156e and its targets. Compared with the wild-type, the *miPEP156e*-overexpressing lines displayed a significantly increased transcript level of miR156e and decreased expression of its target genes *SPL2*, *SPL14,* and *SPL16* (Figure 5C, and 5D). Furthermore, there were no significant phenotypic changes between the transgenic lines and wild-type plants under normal conditions (Figure 5E). However, when exposed to Cd, overexpression of *miPEP156e* significantly enhanced rice resistance to Cd (Figure 5E), which was consistent with the results of miPEP156e external treatment and miR156-OE lines (Figure 3B and 4A). The growth of transgenic lines was barely inhibited by Cd toxicity and exhibited longer root length and higher plant height (Figure 5F). Additionally, the *miPEP156e*-overexpressing lines showed higher biomass in roots and shoots (Figure 5G).

**Figure 5.**
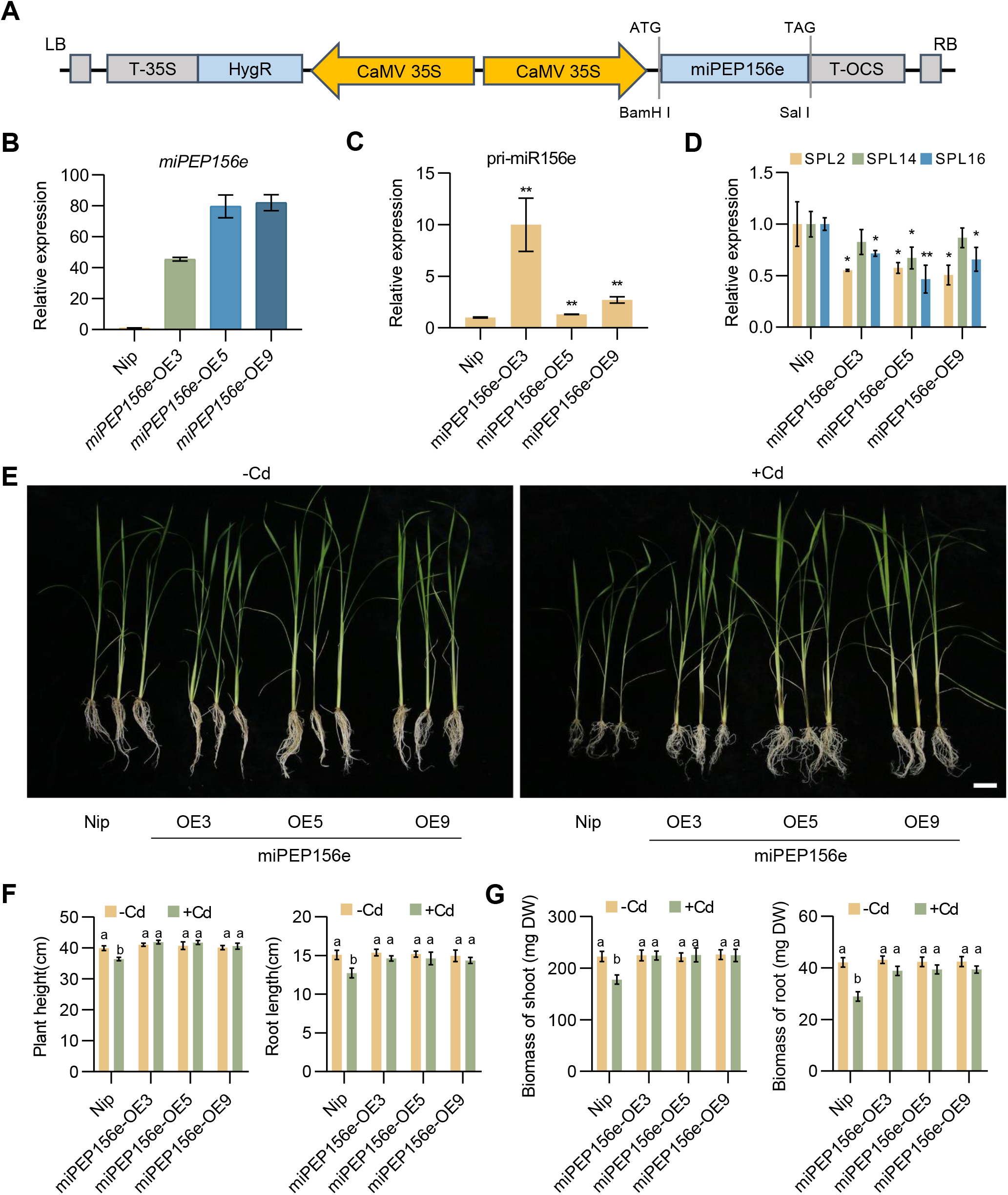
Overexpression of *miPEP156e* regulates miR156e expression and its associated phenotype. A, Schematic representation of the construct used to generate *miPEP156e*-overexpressing lines in Nip. RB, right border; T-OCS, nopaline synthase (NOS) terminator; CaMV 35S, CaMV 35S promoter; HygR, hygromycin resistance gene; T-35S, CaMV 35S terminator; LB, left border. B, Expression level of *miPEP156e* in transgenic rice plants. Error bars indicate SD (n = 3). C, Quantification of pri-miR156e in wild-type and *miPEP156e*-OE seedlings. Error bars indicate SD (n = 3). D, Quantification of miR156e target genes in wild-type and *miPEP156e*-OE seedlings. Error bars indicate SD (n = 3). E, *miPEP156e* overexpression improved the performance of transgenic seedlings under Cd toxicity. F, Plant height and root length of wild-type and *miPEP156e*-OE seedlings after Cd treatment. Error bars indicate SD (n = 8). G, Biomass of shoots and roots of wild-type and *miPEP156e*-OE seedlings after Cd treatment. Error bars indicate SD (n = 8). Asterisks indicate significant differences compared with the corresponding controls (*, *P* < 0,05; **, *P* < 0.01). For F and G, the lowercase letters ‘a’ and ‘b’ indicate significant differences according to Tukey’s test (*P* < 0.05 or *P* < 0.01).

### Development and analysis of *miPEP156* mutants

To further validate the role of miPEP156e in Cd stress tolerance, we generated two *miPEP156e* knockout mutants using CRISPR/Cas9-based genome editing technology. One mutant had a 3-bp deletion at target site 1 and a 1-bp deletion at target site 2 (*miPEP156e-*Cr1), while the other one had a 57-bp deletion (*miPEP156e-*Cr2) (Figure 6A). The homozygous lines were subsequently used for examining the Cd tolerance phenotype. RT-qPCR analysis showed the abundance of pri-miR156e and miR156e significantly decreased in *miPEP156e* knockout mutants (Figure 6B and 6C). In contrast, the expression levels of miR156 targets *SPL2*, *SPL14*, and *SPL16* were significantly up-regulated in the two mutants (Figure 6D), confirming the important role of miPEP156e in regulating the expression of miR156e and its target genes. We also tested the effect of miPEP156e mutation on the Cd tolerance. *miPEP156e-*Cr mutants were more sensitive to Cd stress (Figure 6E), with significantly lower plant height, root length, and biomass than the wild-type (Figure 6F and 6G). These results indicated that miPEP156e is essential for Cd tolerance in rice.

**Figure 6.**
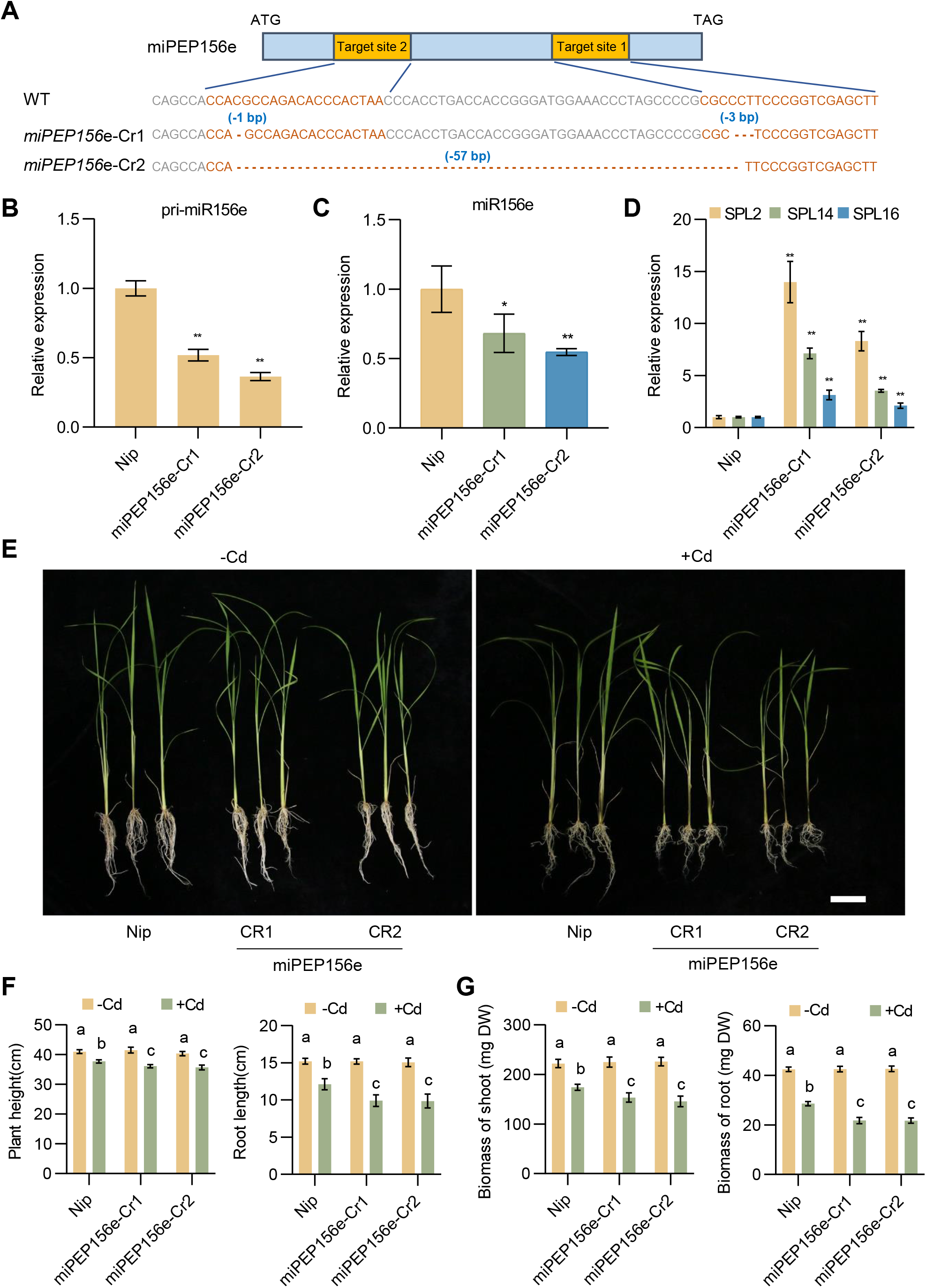
Knockout of *miPEP156e* alters miR156e expression and its associated phenotype. A, Mutation of miPEP156e edited by CRISPR/Cas9 system. Target sequences of two guide RNA (sgRNA) were indicated in orange font. B, Quantification of pri-miR156e in wild-type and *miPEP156e*-Cr seedlings. Error bars indicate SD (n = 3). C, D, Quantification of miR156e (C) and its target genes (D) in wild-type and *miPEP156e*-Cr seedlings. Error bars indicate SD (n = 3). E, Phenotype of in wild-type and *miPEP156e*-Cr seedlings after Cd treatment. F, Plant height and root length of wild-type and *miPEP156e*-Cr after Cd treatment. Error bars indicate SD (n = 8). G, Biomass of shoots and roots of wild-type and *miPEP156e*-Cr seedlings after Cd treatment. Error bars indicate SD (n = 8). Asterisks indicate significant differences compared with the corresponding controls (*, *P* < 0,05; **, *P* < 0.01). For F and G, the lowercase letters ‘a’ to ‘c’ indicate significant differences according to Tukey’s test (*P* < 0.05 or *P* < 0.01).

### miPEP156e diminishes the accumulation of ROS and Cd in rice

The role of miPEP156e in antioxidation and detoxification under Cd stress was analyzed by detecting the contents of ROS and MDA in *miPEP156e*-OE lines and *miPEP156e*-Cr mutants. DAB and NBT staining showed that both the H_2O2 a_nd O^2-^ levels among the three materials were similar in the absence of Cd stress (Figure 7A and 7B). However, upon Cd treatment, the ROS levels were significantly lower in *miPEP156e*-OE plants but higher in *miPEP156e*-Cr mutants than those in the wild-type (Figure 7A and 7B). Consistently, *miPEP156e*-OE plants had lower MDA accumulation levels than *miPEP156e*-Cr mutants and the wild-type (Figure 7C and 7D). Wild-type plants treated with exogenous miPEP156e also accumulated less ROS and MDA, similar to *miPEP156e*-OE plants (Supplemental Figure 5).

**Figure 7.**
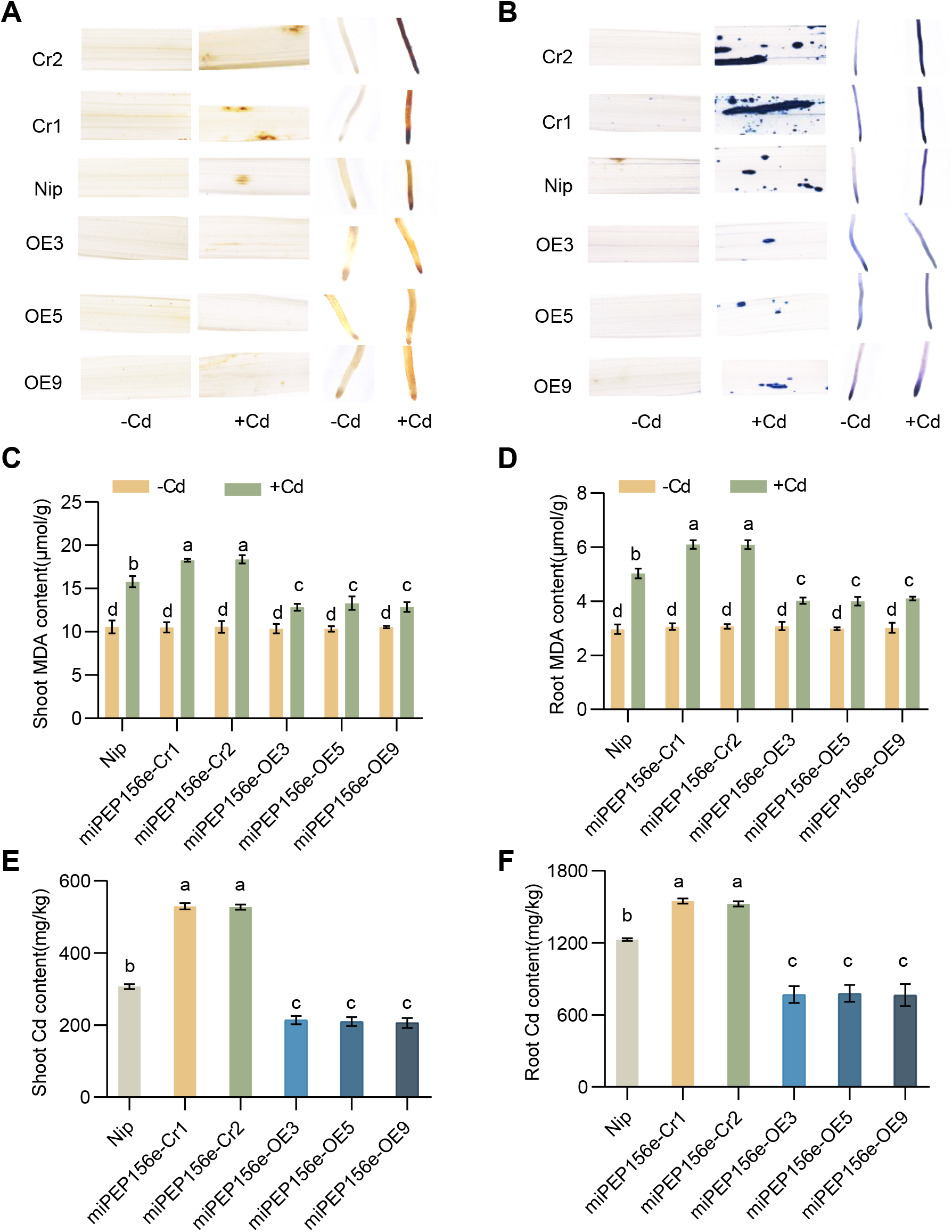
miPEP156e alleviates the accumulation of ROS and Cd in rice under Cd stress. A, DAB staining of leaves and roots of wild-type, *miPEP156e*-OE, and *miPEP156e*-Cr seedlings after water or Cd treatment. The brown color indicates H_2O2 l_evel in each leaf and root. B, NBT staining of leaves and roots of wild-type, *miPEP156e*-OE, and *miPEP156e*-Cr seedlings after water or Cd treatment. The blue color indicates O^2-^ level in each leaf and root. C, Measurement of MDA contents in shoots and roots of wild-type, *miPEP156e*-OE, and *miPEP156e*-Cr seedlings. Error bars indicate SD (n = 3). D, Measurement of Cd contents in shoots and roots of wild-type, *miPEP156e*-OE, and *miPEP156e*-Cr seedlings. Error bars indicate SD (n = 3). For C to F, the lowercase letters ‘a’ to ‘d’ indicate significant differences according to Tukey’s test (*P* < 0.05 or *P* < 0.01).

We further investigated the effect of miPEP156e on Cd content in hydroponically grown plants. As a result, the Cd contents in the shoots and roots of the *miPEP156e*-OE lines were significantly lower than those in wild-type plants, whereas *miPEP156e*-Cr mutants had a much higher concentration of Cd in both shoots and roots, implying that miPEP156e decreased Cd absorption and limited Cd translocation from roots to shoots (Figure 7E and 7F). Lower accumulation of Cd was also observed in the miR156-OE and supplemented-miPEP156e seedlings (Figure 4G and Supplemental Figure 6). These results suggested that miPEP156e modulates the accumulation of miR156 to attenuate the accumulation of ROS and Cd, thus conferring Cd tolerance in rice.

### Transcriptome analysis of downstream pathways modulated by miPEP156e

To better understand the molecular mechanism by which miPEP156e enhances Cd tolerance in rice, we performed transcriptome analysis of roots in *miPEP156e*-OE, *miPEP156e-*Cr, and wild-type Nip seedlings under water and Cd stress conditions. Differentially expressed genes (DEGs) were defined as genes that showed significant expression changes between Cd treatment and mock in three rice materials (Supplemental Figure 7A). Venn diagrams showed that *miPEP156e*-OE9 (Cd) vs *miPEP156e*-OE9 (mock) group and *miPEP156e*-Cr2 (Cd) vs *miPEP156e*-Cr2 (mock) group shared 70% and 64% of the DEGs with Nip (Cd) vs Nip (mock) group, respectively (Figure 8A). Moreover, 3633 overlapping genes were identified by comparing the DEGs in three groups (Figure 8A).

**Figure 8.**
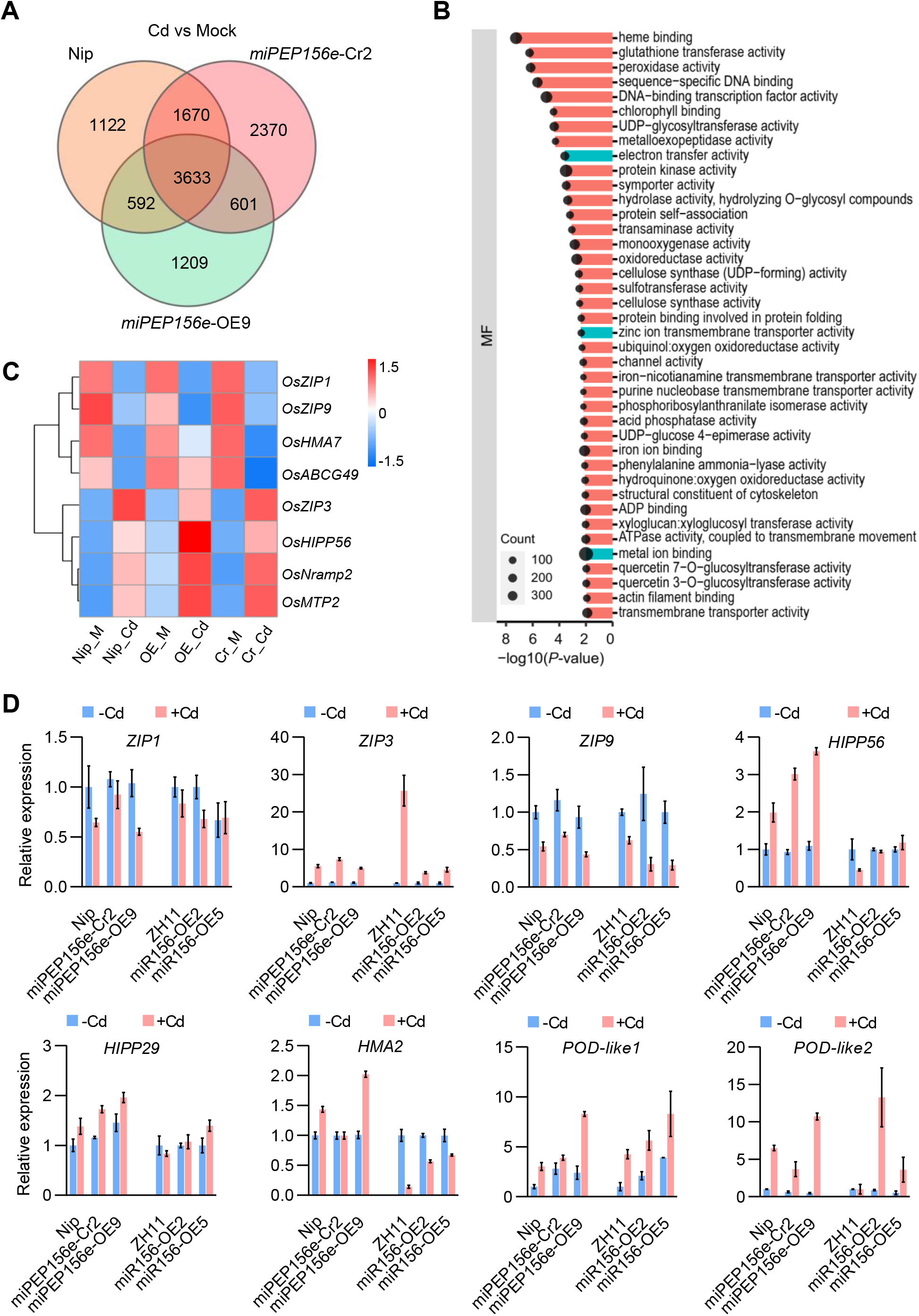
Transcriptomic profiling analysis of *miPEP156e*-OE and *miPEP156e*-Cr in response to Cd stress. A, Venn diagram analyses of differentially expressed genes (DEGs) in Cd-treated wild-type, *miPEP156e*-OE, and *miPEP156e*-Cr root tissues. B Gene ontology (GO) terms enriched in common DEGs shown in A. C Heatmap of DEGs associated with Cd absorption, translocation, and detoxification. D, Expression levels of the Cd-tolerance associated genes in Nip, *miPEP156e*-OE, *miPEP156e*-Cr, ZH11, and miR156-OE seedlings. Error bars indicate SD (n = 3).

To further understand the biological processes and molecular function in which miPEP156e participates, gene ontology (GO) enrichment analysis was performed using the common DEGs across all comparisons. GO enrichment analysis demonstrated that biological processes categories associated with glutathione metabolic process, response to oxidative stress, hydrogen peroxide catabolic process, response to salt stress, and response to hydrogen peroxide, were especially enriched (Supplemental Figure 7B). This was consistent with the conclusion that miPEP156e is involved in the regulation of rice response to Cd stress. More interestingly, molecular function terms related to electron transfer activity, zinc ion transmembrane transporter activity, and metal ion binding, were specifically enriched among the DEGs upon Cd treatment, implicating Cd ion transporters might be involved in miPEP156e-mediated Cd stress response (Figure 8B). Indeed, eight Cd-responsive genes were further identified by comparing the DEGs with the previously reported genes associated with Cd absorption, translocation, and detoxification. Heatmap analysis of these genes further demonstrated that the transcript levels of the genes related to Cd tolerance, including *heavy metal ATPase* (*OsHMA7*), *natural resistance-associated macrophage protein* (*OsNramp2*), *ATP-Binding Cassette G* (*OsABCG49*), *metal tolerance protein gene* (*OsMTP2*), *metallochaperone* (*OsHIPP56*), were up-regulated in *miPEP156e*-OE, while the expression of the genes associated Cd sensitivity, such as *ZRT- and IRT-like protein* (*OsZIP1*, *OsZIP3*, and *OsZIP9*), were down-regulated in *miPEP156e*-OE compared to wild-type Nip (Figure 8C). On the contrary, *miPEP156e*-Cr showed lower expression levels of *OsHMA7* and *OsABCG49* and higher expression of *OsZIP1* than Nip (Figure 8C). In addition, two Cd-resistant genes (*OsHIPP29* and *OsHMA2*) that did not respond to Cd stress showed similar expression profiles in the three rice materials (Supplemental Figure 7C). A comparison analysis was conducted on DEGs with ‘peroxidase activity’, which showed that about 27.9% (17/61) genes related to peroxidase activity were up-regulated in *miPEP156e*-OE (Supplemental Figure 7D and Supplemental Table 2). RT-qPCR analysis of Cd transporter genes and ROS scavenging genes in miR156-OE revealed expression patterns similar to those in *miPEP156e*-OE (Figure 8D). Additionally, we also demonstrated enhanced activities of peroxidase (POD) and superoxide dismutase (SOD) in *miPEP156e*-OE seedlings under Cd toxicity, similar to the results of seedlings treated with exogenous miPEP156e (Supplemental Figure 8). However, *miPEP156e*-Cr mutants exhibited lower SOD and POD activities when exposed to Cd stress compared to the wild-type (Supplemental Figure 8), suggesting that miPEP156e enhanced Cd tolerance in rice by enhancing ROS-scavenging ability and reducing ROS accumulation.

### Identification of novel miPEPs involved in Cd tolerance in rice

Our findings revealed that miR156e-encoded short peptide miPEP156e functions as a positive regulator of Cd tolerance in rice, this inspired us to screen additional miPEPs involved in this progress. The first step for studying miPEPs is to identify the primary transcript of Cd-responsive miRNAs. Previous studies have shown that the first ORF after the transcription start site typically produces a functional miPEP (Lauressergues et al., 2015; Sharma et al., 2020). Based on the potential function of miRNAs and RNA-seq data from NCBI (https://www.ncbi.nlm.nih.gov/), we identified the first ORF of six candidate pri-miRNAs. We then synthesized the six miPEPs corresponding to the six miRNAs, including miR172b, miR396c, miR166b, miR528, miR171c, and miR408.

We next investigated the effects of these miPEPs on the phenotypes of rice seedlings in the presence of Cd toxicity. In our experiments, five of six miPEPs, miPEP172b, miPEP528, miPEP396c, miPEP171c, and miPEP166b, were found to improve the performance of rice seedlings under Cd stress (Figure 9A). After Cd treatment, the seedlings treated with these miPEPs showed longer roots and higher root biomass compared to the control (Figure 9B). We also examined the role of these miPEPs in the expression of their associated miRNAs by analyzing the transcript levels of pri-miRNAs. As expected, they were able to increase the expression of their pri-miRNAs (Figure 9C). In addition, although miPEP408 was found to enhance the transcription of miR408, it was the only small peptide tested that did not affect Cd tolerance in rice (Figure 9).

**Figure 9.**
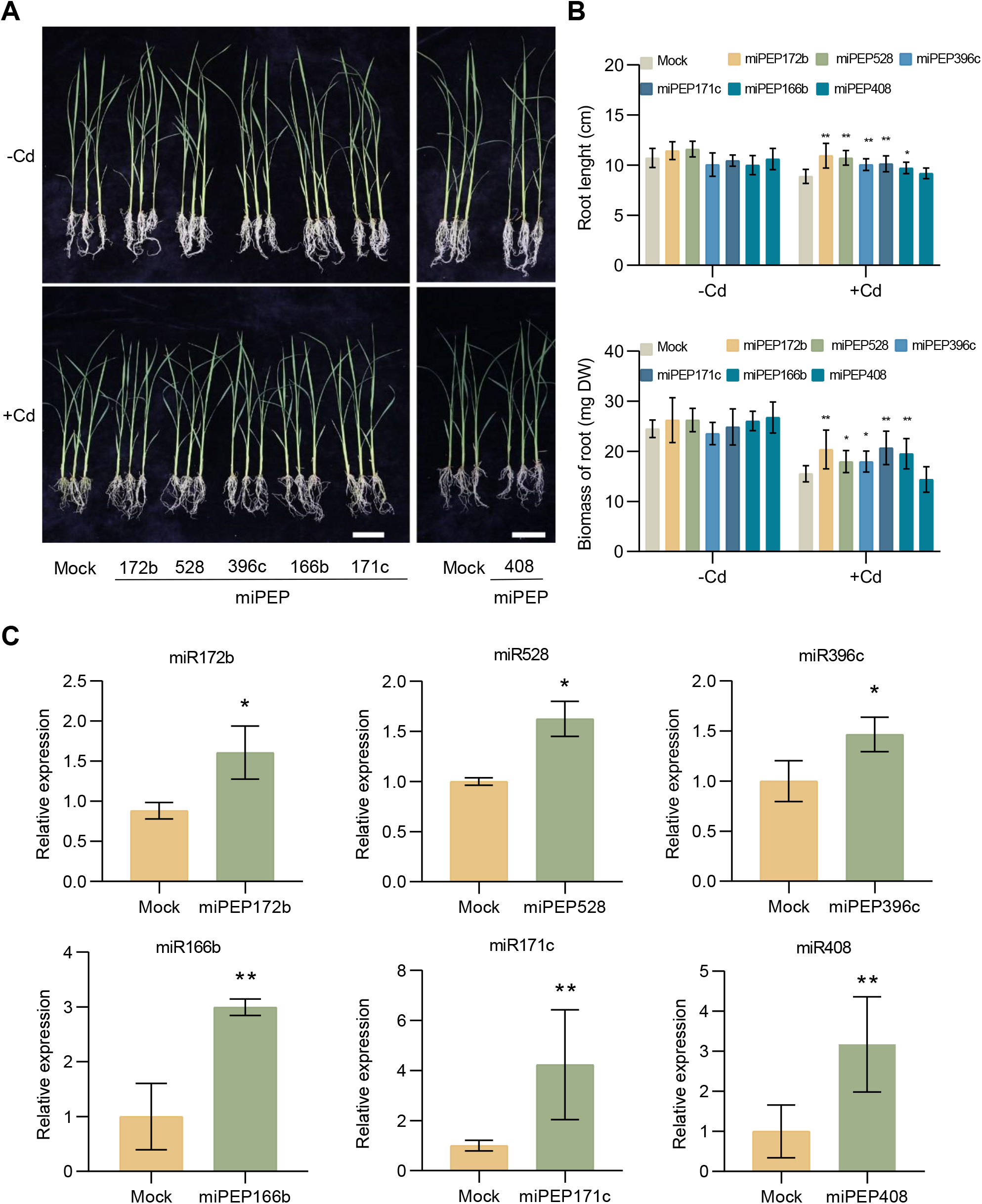
Identification of novel miPEPs involved in Cd tolerance in rice. A, Performance of rice seedlings treated with water or various synthetic miPEPs in the presence of Cd stress. Scale bar = 5 cm. B, Root length and biomass of rice seedlings treated with water or various miPEPs under Cd stress conditions. Error bars indicate SD (n = 8). C, Quantification of pri-miRNAs in the seedlings treated for 6h with water or corresponding miPEPs. Error bars indicate SD (n = 3). Asterisks indicate significant differences compared with the corresponding controls (*, *P* < 0,05; **, *P* < 0.01).

## Discussion

MicroRNAs-encoded regulatory peptides have been recently identified in plants, but their association with abiotic stress tolerance has been rarely reported. In this study, we identified a functional open reading frame encoding a short regulatory peptide, miPEP156e, located in the upstream sequence of pri-miR156e. Further analyses revealed that miPEP156e enhanced the abundance of miR156e, leading to the alleviated accumulation of ROS and Cd under Cd toxicity, and finally conferring Cd tolerance in rice plants. These findings provide new insight into the function of miPEP156e and miR156 in Cd resistance in rice.

Although miR156 has been reported to be responsive to Cd stress, its functional role in relation to Cd stress tolerance in rice remains unknown. In the present study, we revealed that miR156e was induced by Cd stress treatment and showed the highest induction among the miR156 family members (Figure 1). Overexpression of miR156 showed higher tolerance to Cd toxicity, resulting in longer roots and higher plant height (Figure 4). This miR156-mediated Cd tolerance is also conserved in *Arabidopsis*. miR156-overexpressing *Arabidopsis* plants exhibited enhanced tolerance to Cd stress and less Cd in the shoot. However, the knockdown of miR156 led to the increased sensitivity of plants to Cd stress and the higher accumulation of Cd (Zhang et al., 2020). Previous research has demonstrated that miR156 is essential for perceiving and responding to salinity and osmotic stress in both *Arabidopsis* and rice (Cui et al., 2014), indicating that the miR156 regulatory mechanism represents a general adaption strategy for responding to various abiotic stresses.

MicroRNAs, similar to protein-coding genes, are transcribed as long primary transcripts (pri-miRNAs) by RNA polymerase II (Xie et al., 2005). Recent studies have revealed that pri-miRNAs might contain short open reading frames that encode regulatory peptides called miPEPs (Lauressergues et al., 2015; Lauressergues et al., 2022). The presence of miPEPs was revealed in various plant species such as *Arabidopsis thaliana*, *Medicago truncatula*, and *Vitis vinifera*, and even in animals (Lauressergues et al., 2015; Couzigou et al., 2016; Ram et al., 2019; Chen et al., 2020). To determine the functional ORFs present in rice miR156e, the sequence upstream of pre-miR156e was analyzed and four putative ORFs encoding short peptides were identified. Through promoter activity assays, transient expression, and exogenous treatment, we discovered that the peptide (miPEP156e) encoded by the ORF1 could specifically modulate the expression of miR156e and its target *SPLs* genes (Figure 2 and Figure 3). Furthermore, exogenous treatment of plants with miPEP156e induced Cd stress tolerance (Figure 3B), similar to the phenotype observed in miR156-overexpressing plants (Figure 4A), indicating the presence of miPEP in rice. Similarly, the exogenous application of synthetic miPEP858a modulated flavonoid biosynthesis and development in *Arabidopsis* as a result of the enhanced abundance of miR858a (Sharma et al., 2020). The external application of miPEPs from other plant species, such as *Medicago* (miPEP171b), soybean (miPEP172c), and grapevine (miPEP171d1), also led to higher accumulation of their corresponding miRNAs and modulated phenotypes (Lauressergues et al., 2015; Couzigou et al., 2016; Chen et al., 2020), suggesting a common mode of action of miPEPs in plants.

To validate the function of miPEP156e in rice, we generated *miPEP156e*-OE transgenic lines and *miPEP156e*-Cr mutants (Figure 5A and Figure 6A). The *miPEP156e*-OE plants showed a significant increase in the abundance of miR156e, and a decrease in the expression of *SPLs* genes, similar to seedlings treated with exogenous miPEP156e. In contrast, knockout of *miPEP156e* showed a contrasting effect on the gene expression (Figure 5 and Figure 6). Consistently, the *miPEP156e*-OE lines exhibited greater tolerance to Cd stress and better growth compared to wild-type plants (Figure 5). Moreover, we observed reduced accumulation of ROS and MDA in *miPEP156e*-OE lines, as evidenced by the decreased NBT and DAB staining and lower MDA content (Figure 7). On the contrary, *miPEP156e*-Cr mutants showed increased sensitivity toward heavy metal Cd (Figure 6 and Figure 7). These results showed that miPEP156e positively regulates the tolerance to Cd stress by attenuating ROS and MDA production in rice plants. ROS function as important signaling molecules that play important functions in various plant growth and development processes, as well as in the response to environmental cues in plants. On the other hand, the overproduction of ROS during heavy metal stress can cause severe damage to plant tissues (Mittler, 2017; Berni et al., 2019). To counter oxidative stress, plants have developed efficient ROS-scavenging mechanisms that include enzymatic and nonenzymatic antioxidants (Mühling and Läuchli, 2003). Here, we found that the activities of antioxidant enzymes, POD and SOD, were enhanced in *miPEP156e*-OE and seedlings treated with exogenous miPEP156e under Cd stress, while the activation of POD and SOD was restrained in *miPEP156e*-Cr (Supplemental Figure 8). Consistent with this fact, transcriptome analysis revealed that *miPEP156e* overexpression induced a large number of the genes related to peroxidase activity in the presence of Cd toxicity (Supplemental Figure 7), indicating that enhanced ROS-scavenging ability and reduced ROS accumulation contributed to the role of miPEP156e in improving Cd tolerance in rice.

Membrane transporters, including ZIP, HMA, NRAMP, and ABC transporter families, play key roles in heavy metal uptake, transportation, and homeostasis (Takahashi et al., 2012; Xie et al., 2012; Tan et al., 2020). In this study, we discovered that miPEP156e modulates the expression of genes encoding Cd transporters under Cd stress, including *OsZIP1/3/9*, *OsNramp2*, *OsMTP2*, *OsHIPP29/56*, *OsHMA2/7*, *OsABCG49*, and *OsMTP2* (Figure 8C). Consistently, *miPEP156e*-OE lines had substantially lower Cd concentrations in the shoots and roots than *miPEP156e*-Cr mutants and wild-type plants (Figure 7E and 7F). Specifically, we found that *miPEP156e* overexpression down-regulated *OsZIP1*, *OsZIP3*, and *OsZIP9*, which are responsible for Cd uptake, and up-regulated genes contributing to Cd tolerance, such as *OsNramp2*, *OsHIPP29/56*, and *OsHMA2* (Figure 8C and 8D). Differential expression patterns of these genes were observed in *miPEP156e*-Cr (Figure 8C and 8D). The role of miPEP156e in reducing Cd content in the leaves and roots likely resulted from the reduction in Cd absorption and translocation from roots to shoots.

In plants, several secreted peptides encoded by small coding genes have been characterized, known as peptide hormones, which play important roles in plant development and response to biotic and abiotic stress. For example, AtPep3, a hormone-like peptide, is important for conferring both salinity stress tolerance and immune response in *Arabidopsis* (Huffaker et al., 2006; Nakaminami et al., 2018). Moreover, plant elicitor peptide (Pep) signaling helps rice resist insect herbivores and pathogens (Shinya et al., 2018; Shen et al., 2022). Treatment of plants with these peptides can modulate plant growth and stress responses. For instance, 27 CLE peptides were synthesized and roots of plants were treated with them, leading to the identification of CLE25, a small peptide that modulates stomatal control via abscisic acid in long-distance signaling (Takahashi et al., 2018). Since the treatment of plants with synthetic miPEPs can specifically regulate the accumulation of their corresponding mature miRNAs, and produce the comparable phenotypic effects observed in miRNA overexpression plants (Figure 3 and Figure 4). This approach may be useful to analyze the function of novel stress-responsive miRNAs and their regulatory mechanisms. A recent study utilizing this approach demonstrated the role of miPEPs in reducing weed growth (Ormancey et al., 2021). In this study, we identified five novel Cd-tolerance associated miPEPs, including miPEP172b, miPEP528, miPEP396c, miPEP171c, and miPEP166b, which enhanced the expression of their associated miRNAs and improved the performance of rice seedlings under Cd toxicity (Figure 9). The role of miR166b has been characterized (Ding et al., 2018), which suggests that other miPEPs and their corresponding miRNAs may also be involved in the regulation of Cd tolerance in rice.

Taken together, we confirmed the presence of a functional miRNA-encoded peptide miPEP156e and demonstrated its significance in regulating Cd tolerance in rice. We proposed a work model in which miPEP156e modulates the expression of miR156 and its targets by promoting the transcription of pri-miR156e, which improves seedlings’ performance under Cd stress by reducing the accumulation of ROS and Cd in rice plants. Our study also suggested that the exogenous treatment of plants with miPEPs can be a viable alternative to the development of transgenic plants for improving plant tolerance to heavy metals.

## Materials and methods

### Plant materials and growth conditions

Two rice varieties ZH11 (Zhonghua 11) and Nip (Nipponbare) were used as wild-type plants. The miR156-OE transgenic rice plants used in this study were described in a previous study (Liu et al., 2019). The rice seedlings were grown in a greenhouse at 28℃ with a photoperiod of 14 h light/10 h dark. *Nicotiana benthamiana* plants were grown in a greenhouse providing a 14 h photoperiod and constant temperature of 22 °C.

### Rice transformation

To generate the overexpressing *miPEP156e* construct, we cloned the coding sequence of *miPEP156e* into the pCambia1300 vector and introduced it into Nippoonbare (Nip) using *Agrobacterium tumefaciens*-mediated transformation. Homozygous lines were identified using hygromycin screening as described previously (Ma et al., 2013). For generating *miPEP156e* loss-of-function mutant lines, two single-guide RNAs (sgRNAs) were designed online (http://skl.scau.edu.cn/) (Xie et al., 2017) and ligated into the pBUE411 vector (Xing et al., 2014). The resulting construct was introduced into the Nip background, and positive lines were selected in culture solution supplemented with 100 mg L^-1^ hygromycin. Mutations were further confirmed by Sanger sequencing.

### Synthetic peptide assay

The peptides (purity > 95%) were synthesized by Sangon Biotech (Shanghai, China), and dissolved in water at a 5 mM stock concentration. Seedlings were treated with peptide concentrations ranging from 0.1 to 2 μM, which were diluted in water. The sequences of peptides used in this study were listed in Supplemental Table 4.

### Transient expression assay

For the functional ORF analysis, the miR156e promoter regions containing different translation initiation codons were cloned into pCambia2300 vectors containing the *GUS* gene to generate reporter vectors. The constructs were transformed into *A*. *tumefaciens* strain GV3101 and infiltrated into *Nicotiana benthamiana* leaves as described previously (Lu et al., 2021). To determine the activity of four ORFs, the sequences of these reading frames (ORF1-4) were amplified and cloned into pCambia1300 vectors to generate *35S_pro:_ORF1-4* constructs. All the constructs were transfected into rice protoplasts as previously described by the PEG-calcium transfection method (Bart et al., 2006).

### GUS staining

*N*. *benthamiana* leaves harboring the *miR156e_pro:_GUS* (ATG1-4) reporter construct were stained in GUS solution containing 100 mM Na_3P_O_4,_ 10 mM EDTA, 5 mM potassium ferricyanide, 5 mM potassium ferrocyanide, 0.1% Triton X-100, 2 mM X-Gluc (Yisen, Shanghai, China) overnight at 37°C in the dark. The leaves were then washed with an ethanol series of (100%, 95%, 85%, and 75%) to remove chlorophyll and photographed with a Cannon EOS 70D digital camera.

### Measurement of Cd content

Cd content was measured as described previously with some modifications (Sasaki et al., 2014). In brief, the plant samples were dried in an oven at 70 °C for at least 5 days before recording the weight of shoots and roots. Subsequently, 100 mg of the dried shoot and root samples were digested using high-purity HNO_3._ The concentrations of Cd were determined by ICP-OES-Optima 8000 (PerkinElmer, USA).

### Transcriptome analysis

Nip, *miPEP156e*-OE, and *miPEP156e*-Cr rice plants at the four-leaf stage were exposed to 100 μM Cd for 6 h. The roots of each plant were collected with three biological replicates, and total RNA was extracted using the TRNzol Universal Reagent (Tiangen, China). RNA sequencing was performed by Beijing Genomics Institute (BGI, China). After adapter clipping and quality filtering, clean data were remapped to the Nipponbare genome sequence using Bowtie2 software and then quantified using RSEM. DESeq2 software was utilized for analyzing differentially expressed genes (DEGs), which are defined as genes with significant expression changes (|log2(fold change)| > 0.5 and *Q* value < 0.05) in response to Cd treatment compared to mock in all three rice materials.

### Real-time quantitative PCR analysis

Total RNA was extracted using the TRNzol Universal Reagent (Tiangen, China) following the manufacturer’s instructions. First-strand cDNA was synthesized using First-Strand Synthesis Master Mix (Lablead, China). To determine gene expression, RT-qPCR analysis was performed with 2×Realab Green PCR Fast mixture (Lablead, China) in a StepOnePlus real-time PCR instrument (Applied Biosystems). *OsActin* was used as an internal control gene. The miRcute Plant miRNA Isolation Kit (Tiangen, China) was used for the extraction of total miRNAs, and the miRcute plus miRNA First-Strand cDNA Synthesis Kit (Tiangen, China) was used for the reverse transcription of miRcute miRNA first-strand cDNA from the total miRNAs. Osa-U6, a small stably expressed housekeeping RNA molecule, was employed as a quantitative control. All RT-qPCR reactions were performed on three independent replicate samples.

### MDA measurement and ROS examination

Malondialdehyde (MDA) content was measured to quantify lipid peroxidation using the thiobarbituric acid (TBA) method as previously described (Wang et al., 2016). NBT (nitroblue tetrazolium) staining was used for the analysis of superoxide anion (O^2-^) as described before (Jabs et al., 1996). DAB (3, 3’-Diaminobenzidine) staining was used to detect the levels of hydrogen peroxide (H_2O2)_ in rice leaves and roots. DAB staining was performed as described before (Daudi and O’Brien, 2012).

## Author Contributions

LL and XYC performed most of the work and initiated the draft. JMC helped to conduct phenotypic analysis of transgenic rice plants. ZLZ, ZZ, and YYS participated in rice cultivation. YW, SWX, and YNM helped with some experiments and data analysis. RSZ, YYS, and LL conceived the study, obtained funding, and revised the final version of the manuscript. All authors read and approved the final article.

## Supplemental data

**Supplemental Table 1.** Differentially expressed genes in roots in response to Cd treatment.

**Supplemental Table 2.** Gene ontology (GO) analysis of DEGs.

**Supplemental Table 3.** List of peptides used in this study.

**Supplemental Table 4.** List of primers used in this study.

**Supplemental Figure 1.** Analysis of the *MIR156e* transcript.

**Supplemental Figure 2.** Effect of miPEP156e on the phenotypes of rice seedlings cultured in paper pouches under Cd toxicity.

**Supplemental Figure 3.** miPEP156e-ORF2 did not affect miR156e expression and associated phenotype.

**Supplemental Figure 4.** S*P*Ls expression in wild-type and miR156-OE seedlings.

**Supplemental Figure 5.** Exogenous treatment of miPEP156e reduces the accumulation of ROS and MDA in rice under Cd stress.

**Supplemental Figure 6.** Exogenous treatment of miPEP156e reduces Cd accumulation in rice under Cd stress.

**Supplemental Figure 7.** Transcriptomic analysis showing miPEP156e-mediated pathways.

**Supplemental Figure 8.** miPEP156e enhances the activities of POD and SOD in rice seedlings under Cd stress.

