## Supplemental Figure for "miRNA-encoded regulatory peptides modulate cadmium tolerance and accumulation in rice"

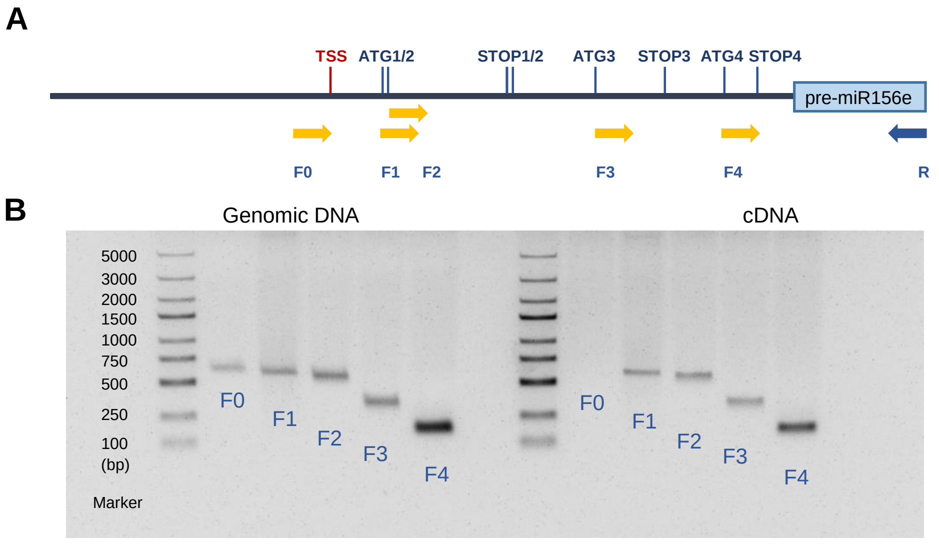


**Supplemental Figure 1. Analysis of the *MIR156e* transcript.**
A, Schematic diagram of primers positions on *MIR156e*. Forward primers are designed in the upstream sequences of pre-miR156e and labeled “F0-F4”. Reverse primer targeted the pre-miR156e sequence was used for PCR amplification. B, PCR analysis of the transcripts of *MIR156e* using different forward primers.

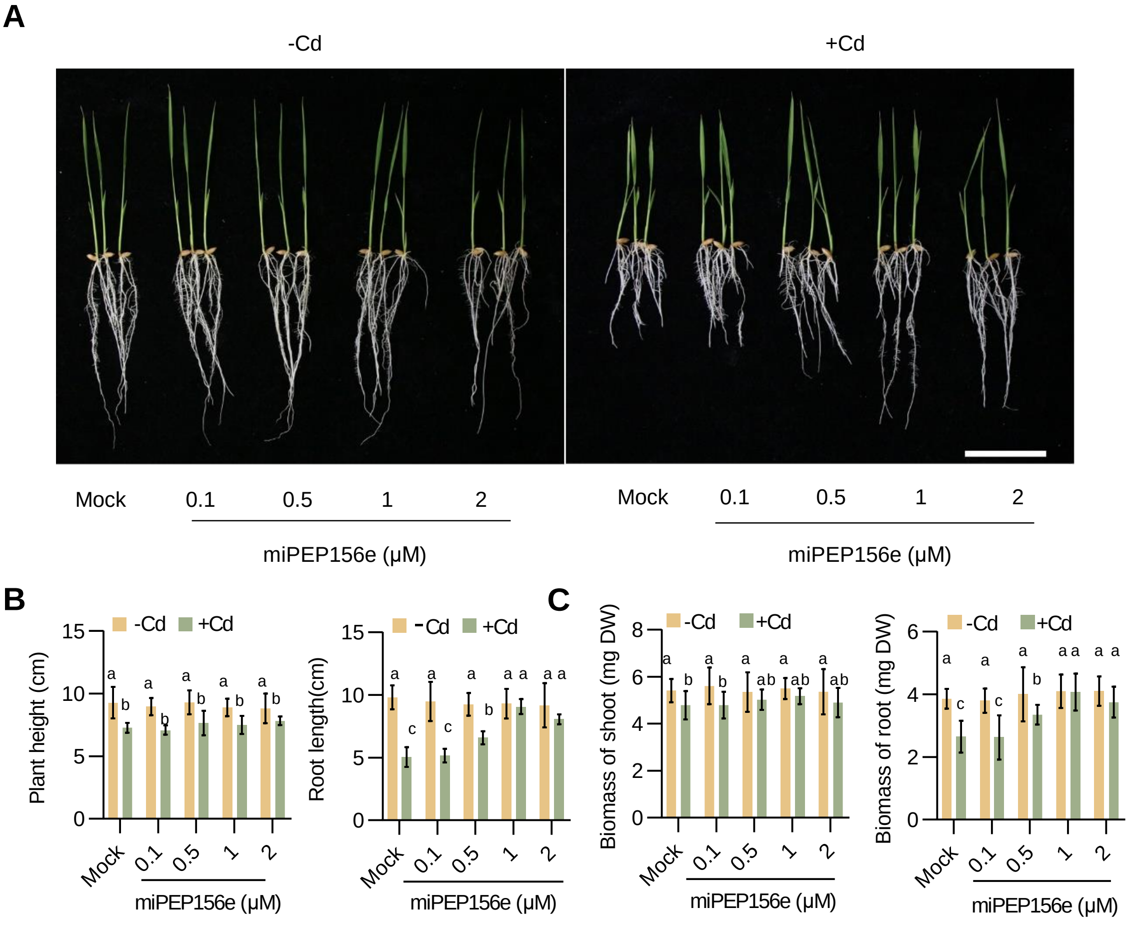


**Supplemental Figure 2. Effect of miPEP156e on the phenotypes of rice seedlings cultured in paper pouches under Cd toxicity.**

A, Performance of 8-day-old seedlings cultured in paper pouches treated with water or different concentrations of miPEP156e in the presence of Cd stress. Scale bar = 5 cm. B, Plant height and root length of 8-day-old seedlings cultured in paper pouches treated with water or different concentrations of miPEP156e under Cd stress conditions. Error bars indicate SD (n = 8). C, Biomass of shoots and roots of 8-day-old seedlings cultured in paper pouches treated with water or different concentrations of miPEP156e under Cd stress conditions. Error bars indicate SD (n = 8). The lowercase ‘a’ to ‘c’ indicate significant differences according to Tukey's test ($P$ < 0.05 or $P$ < 0.01).

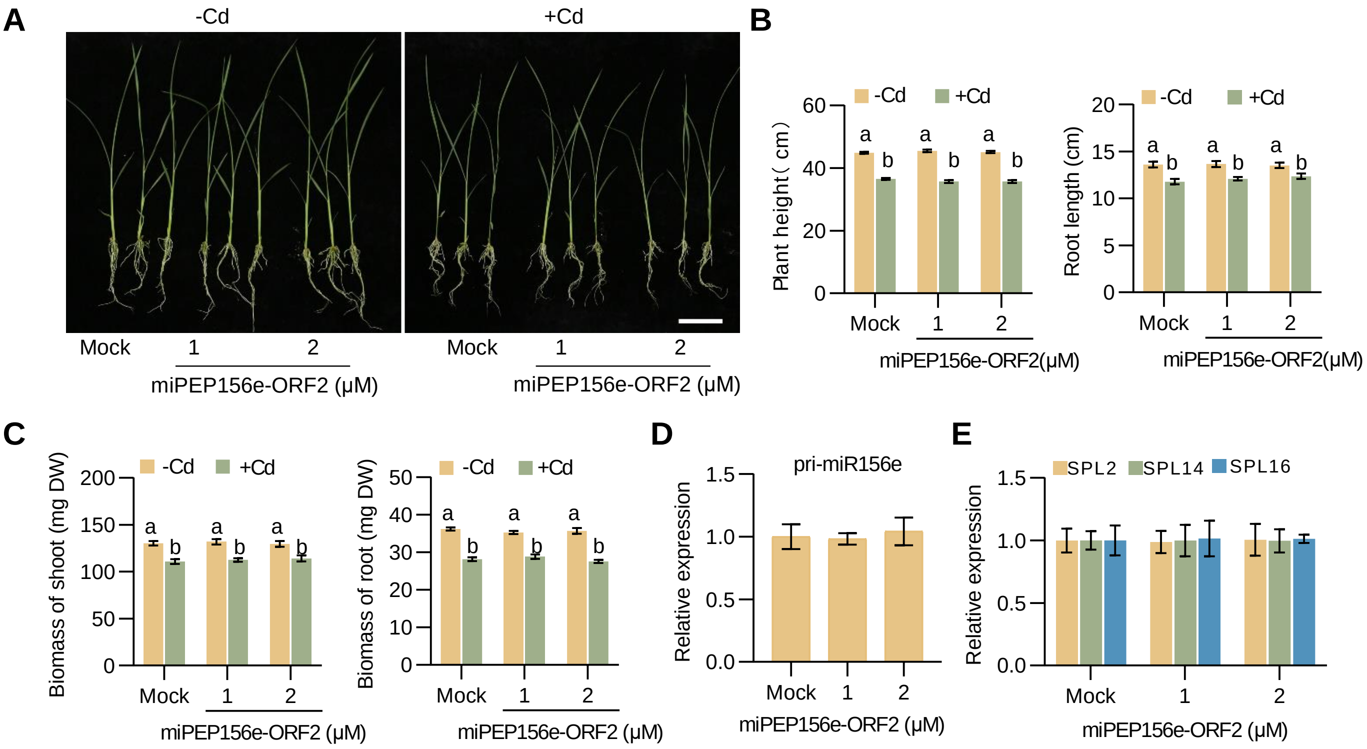


**Supplemental Figure 3. miPEP156e-ORF2 did not affect miR156e expression and associated phenotype.**

A, Performance of 19-day-old seedlings treated with water or different concentrations of miPEP156e-ORF2 in the presence of Cd stress. Scale bar = 5 cm. B, Plant height and root of rice seedlings treated with water or different concentrations of miPEP156e-ORF2 under Cd stress conditions. Error bars indicate SD (n = 8). C, Biomass of shoot and root of rice seedlings treated with water or different concentrations of miPEP156e-ORF2 under Cd stress conditions. Error bars indicate SD (n = 8). D, Quantification of pri-miR156e in the seedlings treated for 6h with water or synthetic miPEP156e-ORF2. Error bars indicate SD (n = 3). E, Quantification of miR156e target genes *SPL2*, *SPL14*, and *SPL16* in the seedlings treated for 6h with water or synthetic miPEP156e-ORF2. Error bars indicate SD (n = 3). The lowercase 'a' and 'b' indicate significant differences according to Tukey's test ($P < 0.05$ or $P < 0.01$).

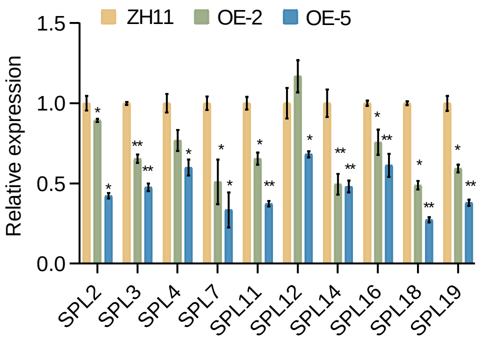


**Supplemental Figure 4. *SPLs* expression in wild-type and miR156-OE seedlings.**
The results are presented as means ± SD of three biological replicates. Asterisks indicate significant differences compared with the corresponding controls (*, $P < 0,05$; **, $P < 0.01$).

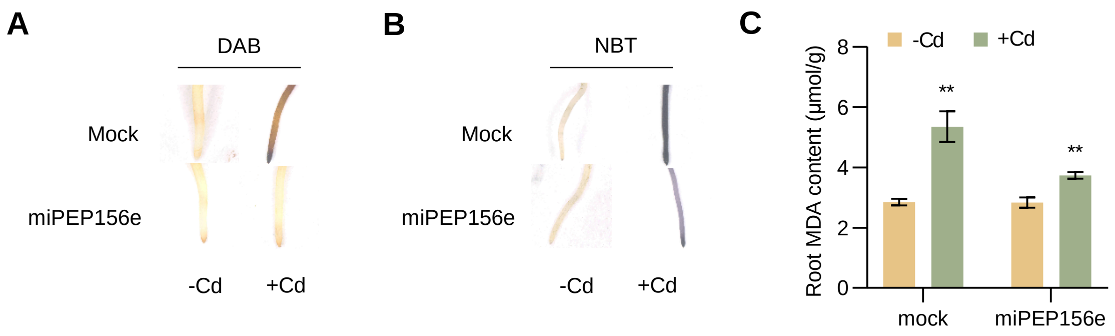


**Supplemental Figure 5. Exogenous treatment of miPEP156e reduces the accumulation of ROS and MDA in rice under Cd stress.**
A, DAB staining of roots of rice seedlings treated with water or miPEP156e under normal or Cd stress conditions. The brown color indicates $H_2O_2$ level in each leaf and root. B, NBT staining of roots of rice seedlings treated with water or miPEP156e under normal or Cd stress conditions. The blue color indicates $O^{2-}$ level in each leaf and root. C, Measurement of MDA contents in roots of rice seedlings treated with water or miPEP156e under normal or Cd stress conditions. Error bars indicate SD (n = 3). Asterisks indicate significant differences compared with the corresponding controls (*, $P < 0,05$; **, $P < 0.01$).

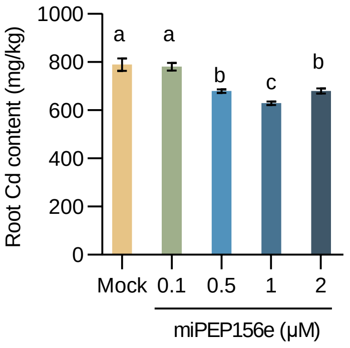


**Supplemental Figure 6. Exogenous treatment of miPEP156e reduces Cd accumulation in rice under Cd stress.**

Measurement of Cd contents in roots of rice seedlings treated with water or miPEP156e under normal or Cd stress conditions. Error bars indicate SD (n = 3). The lowercase ‘a’ to ‘c’ indicate significant differences according to Tukey's test ($P < 0.05$ or $P < 0.01$).

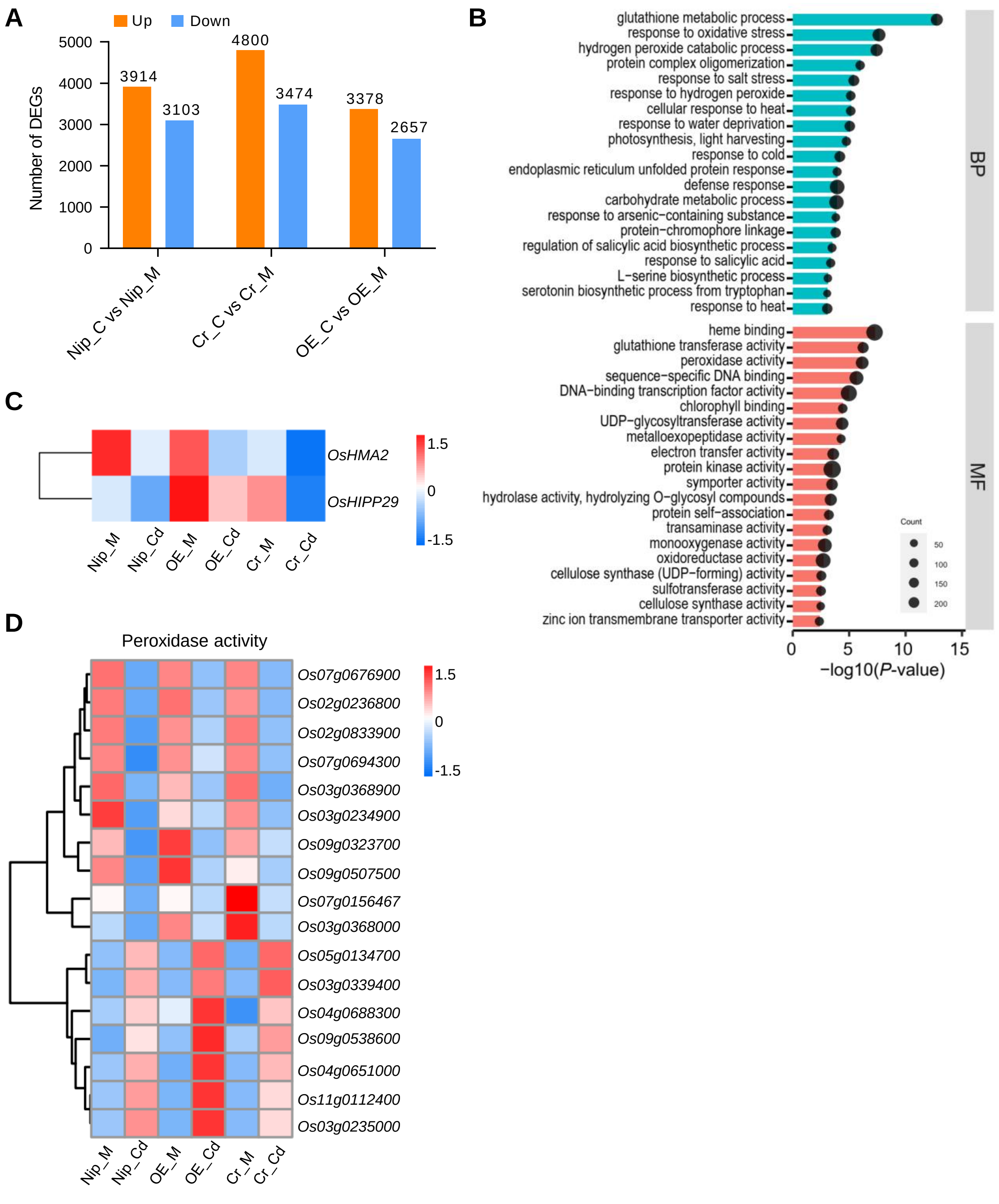


**Supplemental Figure 7. Transcriptomic analysis showing miPEP156e-mediated pathways.**
A, Number of differentially expressed genes (DEGs) between Cd treatment and mock in Nip, *miPEP156e*-OE, and *miPEP156e*-Cr. B Gene ontology (GO) terms enriched in common DEGs shown in Figure 8A. C, Heatmap showing the expression levels of *HMA2* and *HIPP29*. D, Heatmap of DEGs associated with ‘peroxidase activity’.

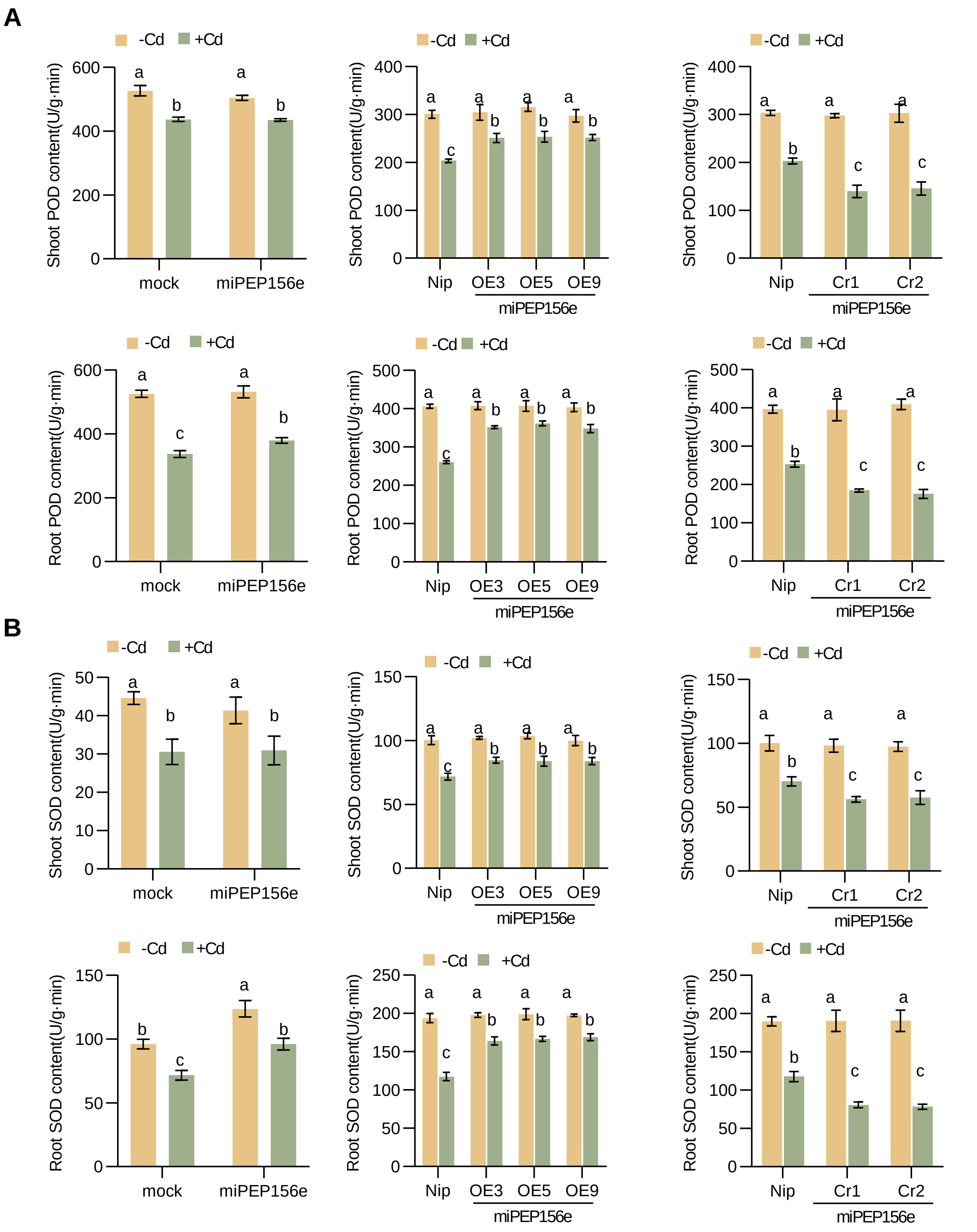


**Supplemental Figure 8. miPEP156e enhances the activities of POD and SOD in rice seedlings under Cd stress.**
A, Measurement of POD activities in the shoots of the wild-type, *miPEP156e*-OE, *miPEP156e*-Cr, and seedlings treated with miPEP156e after water or Cd treatment. Error bars indicate SD (n = 3). B, Measurement of SOD activities in the roots of the wild-type, *miPEP156e*-OE, *miPEP156e*-Cr, and seedlings treated with miPEP156e after water or Cd treatment. Error bars indicate SD (n = 3). The lowercase ‘a’ to ‘c’ indicate significant differences according to Tukey's test ($P < 0.05$ or $P < 0.01$).
